# Gene mutations in members of the PI3K/Akt signalling pathway are related to immune thrombocytopenia pathogenesis

**DOI:** 10.1101/2020.11.16.385963

**Authors:** Rui-Jie Sun, Shu-yan Liu, Xiao-mei Zhang, Jing-jing Zhu, Dai Yuan, Ning-ning Shan

## Abstract

**Purpose:** Immune thrombocytopenic (ITP) is an autoimmune bleeding disease with genetic susceptibility. In this research, we conducted an in-depth genomic analysis of a cohort of patients and elucidated the molecular features associated with the pathogenesis of ITP.

**Method:** High-molecular-weight genomic DNA was extracted from freshly frozen bone marrow blood mononuclear cells (BMBMCs) from 20 active ITP patients. Next, the samples were subjected to molecular genetic analysis by whole-exome sequencing (WES), and the results were confirmed by Sanger sequencing. The signalling pathways and cellular processes associated with the mutated genes were identified with gene ontology (GO) and Kyoto Encyclopedia of Genes and Genomes (KEGG) pathway analyses.

**Results:** The results of this study revealed 3,998 missense mutations involving 2,269 genes in more than 10 individuals. Some unique genetic variants, including phosphatase and tensin homologue (PTEN), insulin receptor (INSR) and coagulation factor C homology (COCH) variants, were the most associated with the pathogenesis of ITP. Functional analysis revealed that these gene mutations mainly affected the phosphoinositide 3 kinase (PI3K)/protein kinase B (Akt) signalling pathways (signal transduction) and platelet activation (immune system).

**Conclusions:** Our findings demonstrate the functional connections between these gene variants and ITP. Although the underlying mechanisms and the impact of these genetic variants remain to be revealed through further investigation, the application of next-generation sequencing in ITP in this paper is valuable for revealing the genetic mechanisms of ITP.

**Summary:** Immune thrombocytopenic (ITP) is an autoimmune bleeding disease with genetic susceptibility. DNA mutation profile of ITP patient bone marrow samples (n=20) were investigated by using next-generation sequencing (NGS), and then confirmed by sanger sequencing method. Our results showed PTEN, INSR and COCH were mutated in all ITP patients. Functional analysis revealed these mutation genes mainly participate PI3K/Akt signaling pathways and platelet activation. These results suggest that genetic alterations might be involved in the pathogenesis of ITP.

## 1. Introduction

Immune thrombocytopenia (ITP) is a complex bleeding disease with autoimmune traits. It is characterized by both decreased platelet production and increased platelet destruction [1]. Patients with ITP present with varying degrees of bleeding tendency, which can cause acute intracranial haemorrhage and life-threatening conditions. Most ITP cases are sporadic, but Rischewski’s group described paediatric ITP cases with a positive family history [2]. Furthermore, genetic susceptibility to ITP has been suggested. One study found that inflammation-related single-nucleotide polymorphisms (SNPs) may be genetic risk factors associated with the disease severity and treatment response of ITP [3]. These results inspired us to use bone marrow blood mononuclear cells (BMBMCs) from a group of primary acute ITP inpatients for whole-exome sequencing (WES) to elucidate the gene variants related to ITP.

The phosphoinositide 3-kinase (PI3K)/protein kinase B (Akt) signalling pathway plays a critical role in regulating the immune response and the release of inflammatory factors in vivo and in vitro by regulating the activation of downstream signalling molecules [4, 5]. In recent years, experimental and clinical evidence has associated perturbations of the PI3K/Akt signal transduction pathway with a number of neoplastic and autoimmune diseases, such as lymphomas[6], chronic and acute lymphocytic leukaemias [7, 8], endometrial cancer [9], bladder cancer [10], rheumatoid arthritis (RA) [11] and ITP [12]. Platelet autophagy is regulated through the PI3K/Akt/mTOR signalling pathway by phosphatase and tensin homologue (PTEN) in ITP. Elevated platelet autophagy may prolong the life span of platelets from ITP patients by inhibiting platelet apoptosis and improving platelet viability [12].

In this study, we identified several genes harbouring an excess number of rare damaging mutations in patients with ITP: PTEN, insulin receptor (INSR) and coagulation factor C homology (COCH). Interestingly, these genes are collectively involved in the signal transduction of the PI3K/Akt signalling pathway and play an important immunomodulatory role in platelet activation. By identifying genetic alterations in ITP patients, our study further enriches the understanding of the pathology of ITP and promotes the identification of potential diagnostic and therapeutic biomarkers for ITP.

## 2. Methods

### 2.1 Patient sample collection and preparation

The study was approved by the Medical Ethical Committee of Shandong Provincial Hospital Affiliated to Shandong University and Shandong Provincial Hospital Affiliated to Shandong First Medical University. Signed informed consent forms were obtained from all participating patients. Twenty newly diagnosed active primary ITP patients, including 12 females and 8 males (age range 17–77 years, median 48 years), seen at the Department of Haematology, Shandong Provincial Hospital, Jinan, China, between May 2017 and November 2018 were enrolled in this study. The diagnosis of ITP was made according to recently published criteria, including patient history, complete blood count, physical examination and peripheral blood smear examination [13]. The platelet counts of patients ranged between 1 and 29 × 10^9^/l, with a median count of 10 × 10^9^/l (Table 1). All the patients required treatment because of clinically significant bleeding. None had been treated with glucocorticosteroids, immunoglobulin or immunosuppressants prior to sampling. Bone marrow blood was collected into heparin-anticoagulant-containing vacutainer tubes. According to the manufacturer’s instructions, mononuclear cells were isolated from heparinized blood by gradient centrifugation with Ficoll-Paque (Pharmacia Diagnostics, Uppsala, Sweden).

**Table 1.** Clinical characteristics of ITP patients

| Sex/Age<br>(year) | Bleeding<br>symptoms | Course of<br>disease<br>(month) | Sex/Age<br>(year) | Bleeding<br>symptoms | Course of<br>disease<br>(month) |
| --- | --- | --- | --- | --- | --- |
| F/19 | PT, EC | 9 | M/59 | NONE | 3 |
| F/43 | EP, GH | 4 | M/63 | PT, GH | 11 |
|  |  |  |  | GH, |  |
| F/38 | PT, GUH | 7 | M/50 | GUH | 21 |
| F/54 | EP | 29 | M/45 | EC, GH | 8 |
| F/43 | PT, GH | 1 | M/59 | NONE | 3 |
| F/32 | PT | 17 | M/63 | PT, GH | 11 |
|  |  |  |  | GH, |  |
| F/48 | PT, GH | 2 | M/50 | GUH | 21 |
| F/69 | EP, GH | 6 | M/45 | EC, GH | 8 |
| F/48 | GH | 12 |  |  |  |
| F/70 | NONE | 14 |  |  |  |
| F/49 | EP, PT | 19 |  |  |  |
| F/17 | EC, EP, GH | 19 |  |  |  |

### 2.2 Targeted exon capture

Genomic DNA was isolated from BMBMCs from 20 active ITP patients using a QIAamp DNA Blood Mini kit (Qiagen, Hilden, Germany) according to the manufacturer’s instructions. Each genomic DNA sample was fragmented using a Covaris LE220 ultrasonicator (Massachusetts, USA) to a fragment size of approximately 200 bp-250 bp. After fragmentation, DNA fragments were end-paired and phosphorylated at the 5’ end and successively adenylated at the 3’ end (following Illumina paired-end protocols), and the libraries ligated to the precapture adaptor were amplified and indexed via PCR. Whole exons were captured with an AI Whole-Exome Enrichment kit (iGeneTech, Beijing, China) after the construction of the sequencing libraries.

### 2.3 Sequencing and sequence alignment

Whole exons were subjected to massive parallel sequencing with 150 paired-end reads on a HiSeqX-Ten sequencer (Illumina, San Diego, California). The program provided with the Illumina Pipeline software package was used to process the raw data in FASTQ format following image analysis and base calling. Clean reads were mapped uniquely for further analysis by removing the adapters and the low-quality reads (defined as those reads for which 50% of reads had a quality value less than 10 and more than 10% Ns in the read length). Filtered reads were successively aligned to the human reference genome sequence (Hg19, NCBI Build 37.5) using the BWA Multi-Vision software package (version 0.7.10).

### 2.4 Variant calling

To ensure accurate variant calling, we applied the recommended best practices for variant analysis in the Genome Analysis Toolkit (GATK, https://www.broadinstitute.org/gatk/guide/best-practices). Base quality score recalibration and insertion and deletion (INDEL) realignment were performed using GATK, with duplicate reads removed by the Picard tools. The sequencing specificity and coverage across each sample were calculated based on the alignments. We employed GATK (v3.3.0) to perform SNP and INDEL discovery and genotyping across all genomic variants. In addition, a strict data analysis quality control (QC) system was used throughout the whole pipeline to guarantee the sequencing data quality.

### 2.5 Variant filtering and annotation

After high-confidence SNPs and INDELs were identified, the SnpEff variant identification tool (http://snpeff.sourceforge.net/SnpEff_manual.html) was employed to (1) verification that the allele frequencies of the mutations in the HapMap, dbSNP, and 1000 Genomes Project databases were ‘0’ and that the allele frequencies of the remaining mutations in the ExAC East Asian AF and ESP6500 AF databases were < 0.1%; (2) verify that the mutations in deleterious coding regions, such as nonsense, missense, frameshift, splice variant and coding INDEL mutations, were retained; (3) perform a cosegregation analysis based on family history using the de novo, autosomal dominant, and autosomal recessive models (excluding mutations that followed other inheritance patterns); and (4) retain those variant predicted to be ‘damaging’ by at least one of the above software packages previously introduced.

### 2.6 Sanger sequencing

Mutations in PTEN, COCH and INSR were confirmed in 20 ITP patients with Sanger sequencing. The primers used to amplify the exon region by PCR are shown in Table 2 and Supplemental S2.xls. Sequencing data were obtained by Beijing Genomics Institute (BGI, Shenzhen, China) and analysed using SeqMan Lasergene software. The resulting sequences were compared with the published sequences of PTEN (GenBank accession number NM_005960 and corresponding protein sequence NP_005951.1), COCH (GenBank accession number NM_002458.3 and corresponding protein sequence NP_002449.2), and INSR (GenBank accession number NM_005961.3 and corresponding protein sequence NP_005952.2).

**Table 2.**
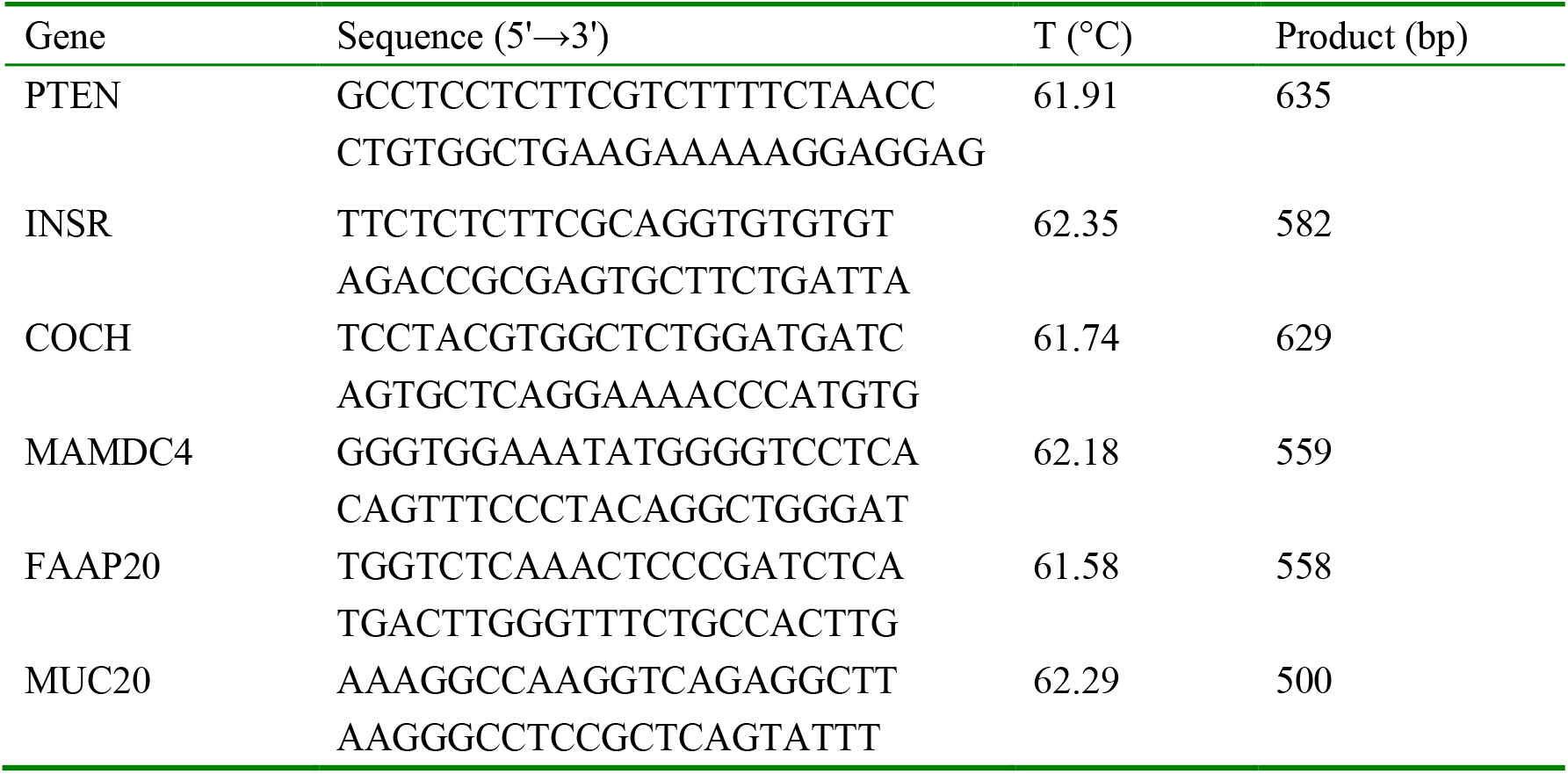
Primers and conditions for the Sanger sequencing in this study

### 2.7 Statistics analysis

Statistical analysis was performed using SPSS version 16.0 software (SPSS, Chicago, IL, USA). Differences between groups were compared using one-way ANOVA. A p-value<0.05 was considered significant.

## 3 Results

### 3.1 Functional analysis of the missense mutation-containing genes

To identify genomic alterations associated with the pathogenesis of ITP, we used WES to detect the DNA mutation profiles of BMBMCs from ITP patients (n=20). A total of 3,998 missense mutations involving 2,269 genes were identified in more than 10 individuals (Supplemental S1.xls). Next, the potential functions of the mutated genes were analysed using Kyoto Encyclopedia of Genes and Genomes (KEGG) and gene ontology (GO) pathway analyses. The functional analysis revealed that most of the genes were related to signal transduction. The biological processes, cellular components and molecular functions significantly associated with the mutated genes were obtained from GO analysis (Fig 1A). In the biological process category, the mutated genes were highly enriched in cellular processes and biological processes. In the cellular component category, the mutated genes were mainly associated with the terms cell, organelle and membrane. In the molecular function category, the mutated genes were highly associated with binding, signalling and molecular transducer activity.

**Figure 1.**
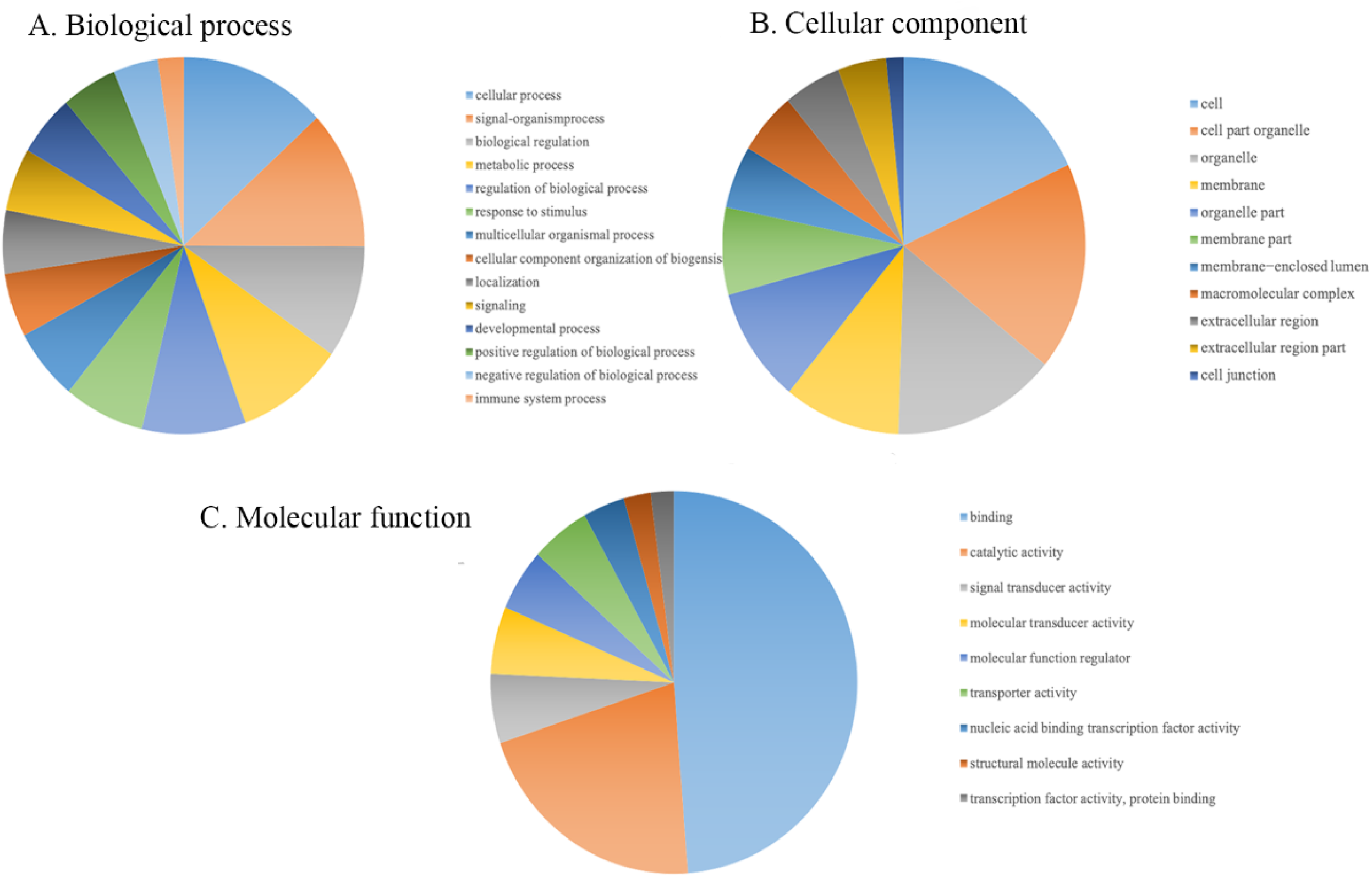
GO Analysis. **A.** Biological process; **B.** Cellular component; **C.** Molecular function.

Furthermore, KEGG analysis demonstrated that the mutated genes were collectively associated with tight junctions, regulation of the actin cytoskeleton, the Rap1 signalling pathway, and focal adhesion and cell adhesion molecules (CAMs) (Fig 1B).

### 3.2 Identification of genes mutated in all analysed ITP patients

The results indicated that four genes (PTEN, INSR, COCH and MAM domain-containing 4 (MAMDC4)) harboured missense mutations in all ITP patients (Supplemental S1.xls). These genes (PTEN, INSR, COCH, and MAMDC4) are involved in the PI3K/Akt signalling pathway and thus might affect the activation of platelets and be associated with the pathogenesis of ITP. In addition, the PTEN gene regulates autophagy via the mTOR signalling pathway to mediate the onset of ITP. In addition, INSR is involved in HIF-1 pathway regulation, and COCH is involved in the regulation of platelet activation during the immune response.

Four pathways (the HIF-1, mTOR, and PI3K/Akt signalling pathways (related to signal transduction) and the platelet activation pathway (related to the immune system)) showed the highest association with ITP in this study, and the details of these pathways are shown in Table 1 (Supplementary Table S2.xls).

In addition, we found Fanconi anaemia-associated protein, 20 kDa (FAAP20) mutations in DNA samples from 19 patients and mucin 20 (MUC20) mutations in DNA samples from 18 patients. The FAAP20 and MUC20 proteins are also involved in platelet activation and the regulation of the PI3K/Akt signalling pathway, suggesting that they may also affect the pathogenesis of ITP.

### 3.3 Sanger sequencing analysis

Moreover, the mutations in the PTEN, INSR, COCH, MAMDC4, FAAP20 and MUC20 genes were also verified with Sanger sequencing. The results suggested that genetic alteration of genes might be associated with the pathogenesis of ITP. Fig 2A–2F shows a novel missense mutation in each mutated protein.

**Figure 2.**
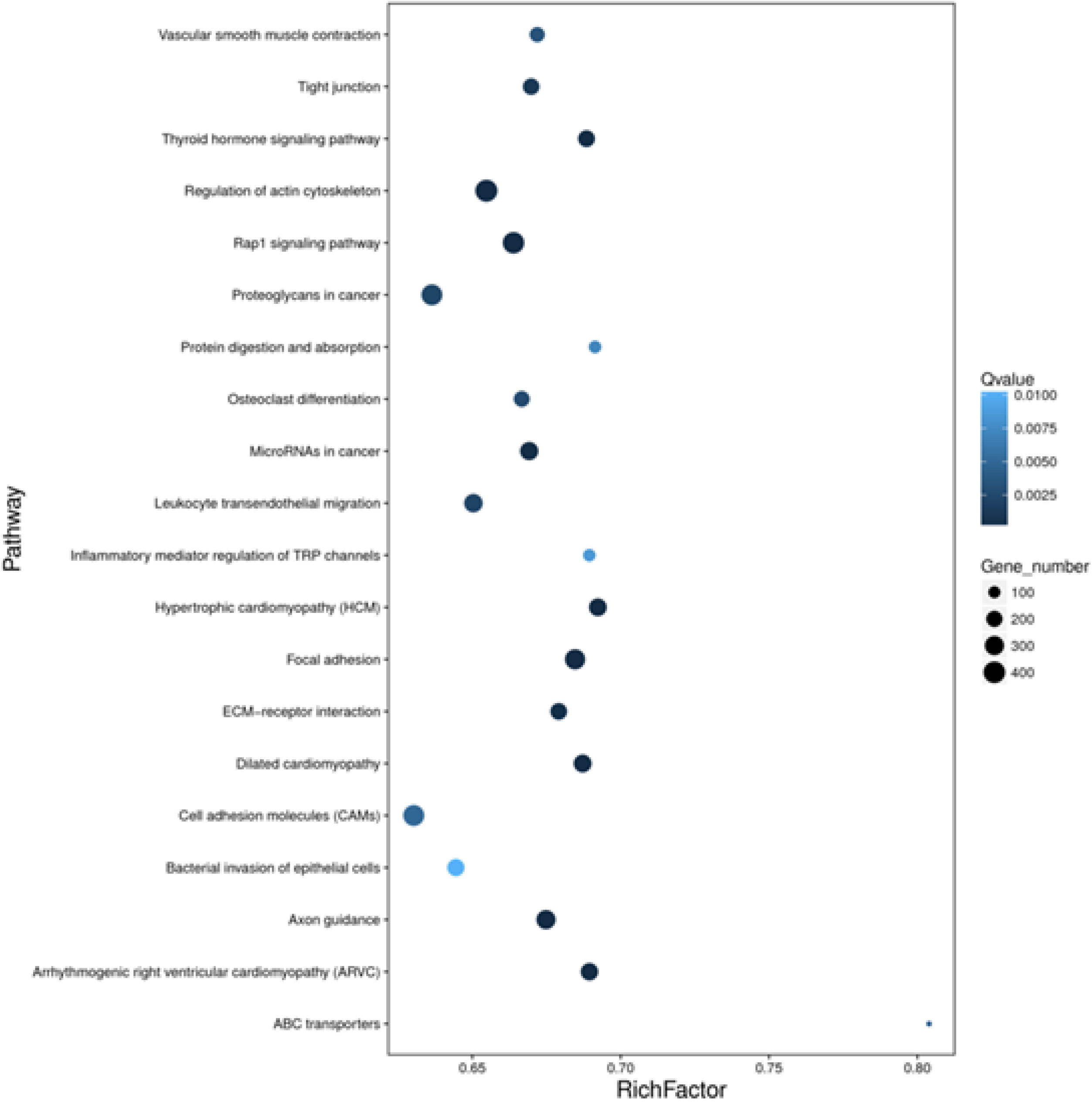
KEGG Pathway. The p-values obtained by using the Fisher’s exact test showed the functional classifications and pathways in differentially expressed protein, which are displayed in a bubble chart.

**Figure 3.**
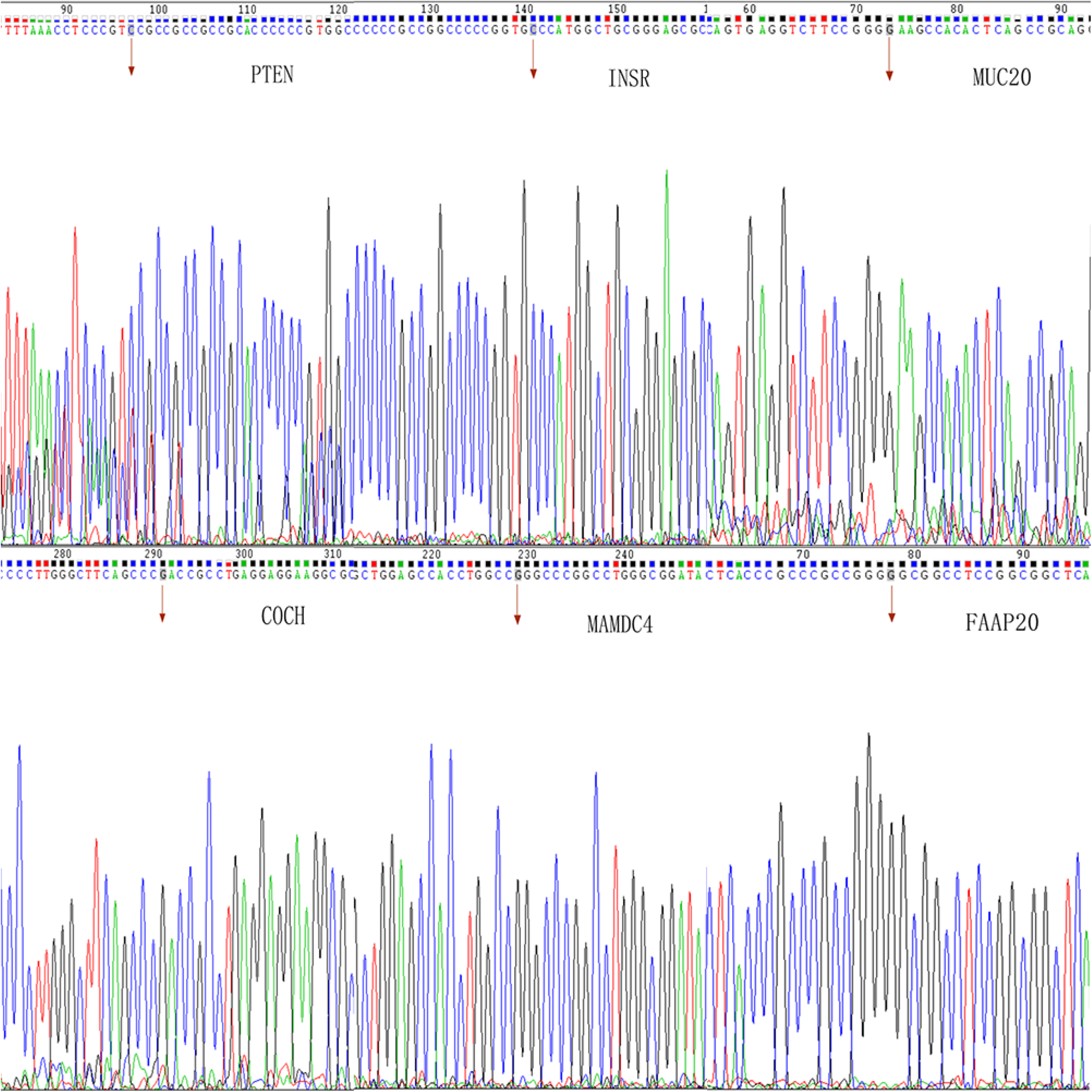
Mutated Gene. **A.** PTEN; **B.** INSR; **C.** COCH; **D.** MAMDC4; **E.** FAAP20; **F.** MUC20.

## 4. Discussion

ITP is a complex disease featuring autoimmune bleeding that is affected by multiple genetic and environmental influences [3, 14, 15]. In the plasma of ITP patients, platelet membrane proteins become antigenic and then stimulate the immune system to produce antibodies, eventually resulting in T cell immune imbalance and thrombocytopenia [1]. Several DNA SNPs play an important role in the pathogenesis of ITP [16, 17]. Rischewski’s group proposed the existence of genetic susceptibility to ITP by describing paediatric ITP cases with a positive family history [2]. In this study, we found several DNA missense mutations related to the PI3K/Akt signalling pathway in BMBMCs from ITP patients, which may indicate that this pathway is involved in the pathogenesis of ITP.

Our previous quantitative proteomics analysis showed that apoptosis-related proteins (HSPA8, HSPA6, ITGB3, YWHAH, and PRDX6) [18] and autophagy-related proteins (HSPA8, PARK7, YWHAH, ITGB3 and CSF1R) were significantly abnormally expressed in ITP BMBMC samples compared to normal controls. We found that these differentially expressed proteins were significantly downregulated using parallel reaction monitoring (PRM) verification, except for CSF1R, which was upregulated [18]. KEGG enrichment analysis showed that these differentially expressed proteins were also closely related to the PI3K/Akt signalling pathway [18].

The PI3K pathway is an essential pathway for various cellular processes, and it is also one of the most frequently activated signal transduction pathways in human cancer and autoimmune disease. The central role of Akt in the PI3K pathway makes it one of the most activated downstream effectors [17]. Akt interacts with the cytoplasmic domain of GPIbα [19] and transduces signalling in response to the vWF-GPIbα interaction, leading to platelet activation [20]. PI3K/Akt signalling may be antagonized by the tumour suppressor PTEN, which was identified as a frequently mutated gene in many types of tumours, particularly endometrial, skin, brain, and prostate tumours [21, 22]. Our previous research showed that perturbations in normal autophagy, which may be caused by deletion of autophagy-related genes such as ATG7 and abnormal signalling due to overexpression of mTOR, lead to abnormal platelet and megakaryocyte functions [23]. mTOR is a key kinase and negative regulator of the PI3K/Akt/mTOR signalling pathway and can regulate cell proliferation, growth, survival, and angiogenesis under physiological conditions and in the presence of environmental stress [24]. PTEN is a key positive regulatory molecule of autophagy that blocks the inhibitory effect of PI3K/PKB on autophagy, thereby activating autophagy and inducing autophagosome formation [25]. In experiments in vitro, indirubin was observed to restore the expression of programmed cell-death 1 (PD1) and PTEN in the CD4+ T cells of ITP patients, leading to subsequent attenuation of Akt/mTOR pathway signalling and modulation of T cell homeostasis [26]. Thus, it may be hypothesized that PTEN mutations lead to activation of the PI3K/Akt/mTOR pathway and inhibition of autophagy and play a role in ITP initiation and progression.

INSR is the central mediator in the insulin response upstream of PI3K that induces tyrosine phosphorylation of INSR substrates and subsequent activation of enzymes downstream of PI3K [27, 28]. Several studies have shown that PI3K/Akt pathway activation can be induced by insulin and that insulin acts as an indispensable effector [29, 30]. As a downstream molecule of the PI3K/Akt pathway, mTOR not only influences autophagy balance but also increases hypoxia-inducible factor 1α (HIF-1α) activity and the production of reactive oxygen species (ROS), leading to oxidative stress in cells [31]. Caroline et al. have shown that insulin regulates HIF-1 subunit accumulation and activation through a PI3K/mTOR-dependent pathway, resulting in increased vascular endothelial growth factor (VEGF) expression [32].

VEGF is a key angiogenic factor involved in a wide variety of biological processes, including embryonic development, tumour progression and metastasis, and is regulated by platelet-derived growth factor, insulin, insulin-like growth factor-I, and tumour necrosis factor [33, 34]. Functional analysis in this study revealed that mutated INSR is involved in the PI3K/Akt signalling pathway and HIF-1 signalling pathway in ITP patients. Although the in-depth mechanism underlying the effects of INSR mutation on ITP pathogenesis remains to be uncovered, INSR and PTEN exon mutations may be involved in the PI3K/Akt signalling pathway, further affecting the expression of downstream molecules and eventually participating in the pathogenesis of ITP.

In addition, functional clustering analysis showed that COCH participates in platelet activation. The COCH gene was the first gene identified to cause vestibular dysfunction [35]. COCH encodes cochlin, which contains a short signal peptide (SP), an N-terminal factor C homology (FCH or LCCL) domain and two von Willebrand factor A-like domains (vWFA1 and vWFA2) [35, 36]. vWFA domains are known for their ability to induce self-aggregation in response to shear stress and adherence to macrophages, platelets or leukocytes [36]. PI3K association with the cytoplasmic domain of GPIbα transduces vWF-binding signalling, leading to Akt activation [20, 37]. Some patients with DFNA9 (a vestibular disorder) develop vascular diseases such as cerebral ischaemia and acute myocardial infarction, and vWFA domains have been implicated in increased shear-induced platelet aggregation (SIPA) [36]. However, the function of the COCH gene in ITP pathogenesis remains to be fully elucidated. The MAMDC4 protein is associated with a unique endocytic mechanism observed in the intestine of mammals [38], which may be related to the autophagy activities mediated by the PI3K/Akt signalling pathway. In addition, FAAP20 and MUC20 have also been shown to participate in the PI3K/Akt pathway and platelet activation in most ITP samples. Further studies are needed to improve the understanding of the role of missense mutations and related functional pathways in ITP.

Wang et al. experimentally enhanced autophagy-related protein and autophagic flux in the PI3K/Akt/mTOR signalling pathway, inhibiting apoptosis and improving platelet viability, to alleviate platelet destruction and prolong the life span of platelets from ITP patients [12]. Furthermore, microRNAs act by targeting insulin-like growth factor 2 mRNA-binding protein 1 (IGF2BP1), and the subsequent downregulation of insulin-like growth factor 2 (IGF-2) causes inhibition of the PI3K/Akt pathway, which is involved in the mesenchymal stem cell (MSC) deficiency seen in ITP [39]. We previously identified abnormal expression of multiple proteins in the PI3K/Akt pathway in patient groups compared with control groups via protein profiling analysis [18]. In support of these finding, this study confirmed the presence of mutation in the exons of genes encoding proteins in the PI3K/Akt pathway (PTEN, INSR, COCH, MAMDC4, FAAP20 and MUC20) in ITP bone marrow samples, further verifying the important role of this signalling pathway in ITP pathogenesis. However, little is known about the detailed transcription processes and pathological effects of mutated proteins leading to thrombocytopenia.

In conclusion, our findings improve the understanding of the PI3K/Akt signalling pathway and, more significantly, suggest therapeutic targets and research directions for ITP caused by specific gene mutations or other pathogenic factors. Future work is needed to determine how the transcription and translation of these mutated genes in the PI3K/Akt pathway affect the occurrence and development of ITP.

## Ethical approval and consent to participate

Informed consent was obtained from each participating patient and/or legal guardian. Ethical approval for the study was obtained from the Medical Ethical Committee of Shandong Provincial Hospital Affiliated to Shandong University and Shandong Provincial Hospital Affiliated to Shandong First Medical University.

## Competing interests

The authors declare no competing interests.

## Funding

The work was supported by grants from by Taishan Youth Scholar Foundation of Shandong Province (tsqn201812140), Academic promotion programe of Shandong First Medical University (2019RC018), Taishan Scholar Foundation of Shandong Province, National Natural Science Foundation of China (No. 81570104), Key Research and Development Project of Jinan (No. 201907021; No. 201907026), Key Research and Development Program of Shandong Province (No. 2018CXGC1213); Technology Development Projects of Shandong Province (No. 2017GSF18189).

## Reference

1. Shan, N.N., et al., Targeting autophagy as a potential therapeutic approach for immune thrombocytopenia therapy. Crit Rev Oncol Hematol, 2016. 100: p. 11–5.

2. Rischewski, J.R., et al., Idiopathic thrombocytopenic purpura (ITP): is there a genetic predisposition? Pediatr Blood Cancer, 2006. 47(5 Suppl): p. 678–80.

3. Li, J., et al., Inflammation-Related Gene Polymorphisms Associated With Primary Immune Thrombocytopenia. Front Immunol, 2017. 8: p. 744.

4. Xu, P., et al., Regulatory roles of the PI3K/Akt signaling pathway in rats with severe acute pancreatitis. PLoS One, 2013. 8(11): p. e81767.

5. Cianciulli, A., et al., PI3k/Akt signalling pathway plays a crucial role in the anti-inflammatory effects of curcumin in LPS-activated microglia. Int Immunopharmacol, 2016. 36: p. 282–290.

6. Cannon, M.L. and E. Cesarman, The KSHV G protein-coupled receptor signals via multiple pathways to induce transcription factor activation in primary effusion lymphoma cells. Oncogene, 2004. 23(2): p. 514–23.

7. Schade, A.E., M.W. Wlodarski, and J.P. Maciejewski, Pathophysiology defined by altered signal transduction pathways: the role of JAK-STAT and PI3K signaling in leukemic large granular lymphocytes. Cell Cycle, 2006. 5(22): p. 2571–4.

8. Perera, Y., et al., Clinical-Grade Peptide-Based Inhibition of CK2 Blocks Viability and Proliferation of T-ALL Cells and Counteracts IL-7 Stimulation and Stromal Support. Cancers (Basel), 2020. 12(6).

9. Zheng, Y., et al., HDAC6, modulated by miR-206, promotes endometrial cancer progression through the PTEN/AKT/mTOR pathway. Sci Rep, 2020. 10(1): p. 3576.

10. Egawa, H., et al., The miR-130 family promotes cell migration and invasion in bladder cancer through FAK and Akt phosphorylation by regulating PTEN. Sci Rep, 2016. 6: p. 20574.

11. Feng, F.B. and H.Y. Qiu, Effects of Artesunate on chondrocyte proliferation, apoptosis and autophagy through the PI3K/AKT/mTOR signaling pathway in rat models with rheumatoid arthritis. Biomed Pharmacother, 2018. 102: p. 1209–1220.

12. Wang, C.Y., et al., Enhancing autophagy protects platelets in immune thrombocytopenia patients. Ann Transl Med, 2019. 7(7): p. 134.

13. Provan, D., et al., International consensus report on the investigation and management of primary immune thrombocytopenia. Blood, 2010. 115(2): p. 168–86.

14. Vandrovcova, J., et al., FAS mutations are an uncommon cause of immune thrombocytopenia in children and adults without additional features of immunodeficiency. Br J Haematol, 2019. 186(6): p. e163–e165.

15. Heelan, B.T., et al., Effect of anti-CD20 (rituximab) on resistant thrombocytopenia in autoimmune lymphoproliferative syndrome. Br J Haematol, 2002. 118(4): p. 1078–81.

16. Bergmann, A.K., R.F. Grace, and E.J. Neufeld, Genetic studies in pediatric ITP: outlook, feasibility, and requirements. Ann Hematol, 2010. 89 Suppl 1: p. S95–103.

17. Martini, M., et al., PI3K/AKT signaling pathway and cancer: an updated review. Ann Med, 2014. 46(6): p. 372–83.

18. Liu, S.Y., et al., Significant reductions in apoptosis-related proteins (HSPA6, HSPA8, ITGB3, YWHAH, and PRDX6) are involved in immune thrombocytopenia. J Thromb Thrombolysis, 2020.

19. Mu, F.T., et al., A functional 14-3-3zeta-independent association of PI3-kinase with glycoprotein Ib alpha, the major ligand-binding subunit of the platelet glycoprotein Ib-IX-V complex. Blood, 2008. 111(9): p. 4580–7.

20. Yin, H., et al., The role of Akt in the signaling pathway of the glycoprotein Ib-IX induced platelet activation. Blood, 2008. 111(2): p. 658–65.

21. Li, J., et al., PTEN, a putative protein tyrosine phosphatase gene mutated in human brain, breast, and prostate cancer. Science, 1997. 275(5308): p. 1943–7.

22. Ali, I.U., L.M. Schriml, and M. Dean, Mutational spectra of PTEN/MMAC1 gene: a tumor suppressor with lipid phosphatase activity. J Natl Cancer Inst, 1999. 91(22): p. 1922–32.

23. Sun, R.J. and N.N. Shan, Megakaryocytic dysfunction in immune thrombocytopenia is linked to autophagy. Cancer Cell Int, 2019. 19: p. 59.

24. Kim, Y.C. and K.L. Guan, mTOR: a pharmacologic target for autophagy regulation. J Clin Invest, 2015. 125(1): p. 25–32.

25. Guo, Y., et al., Thymosin alpha 1 suppresses proliferation and induces apoptosis in breast cancer cells through PTEN-mediated inhibition of PI3K/Akt/mTOR signaling pathway. Apoptosis, 2015. 20(8): p. 1109–21.

26. Zhao, Y., et al., Indirubin modulates CD4(+) T-cell homeostasis via PD1/PTEN/AKT signalling pathway in immune thrombocytopenia. J Cell Mol Med, 2019. 23(3): p. 1885–1898.

27. Saltiel, A.R. and J.E. Pessin, Insulin signaling pathways in time and space. Trends Cell Biol, 2002. 12(2): p. 65–71.

28. Hemi, R., et al., p38 mitogen-activated protein kinase-dependent transactivation of ErbB receptor family: a novel common mechanism for stress-induced IRS-1 serine phosphorylation and insulin resistance. Diabetes, 2011. 60(4): p. 1134–45.

29. Zhou, M., et al., Effect of central JAZF1 on glucose production is regulated by the PI3K-Akt-AMPK pathway. FASEB J, 2020. 34(5): p. 7058–7074.

30. Mazibuko-Mbeje, S.E., et al., Aspalathin-Enriched Green Rooibos Extract Reduces Hepatic Insulin Resistance by Modulating PI3K/AKT and AMPK Pathways. Int J Mol Sci, 2019. 20(3).

31. Nacarelli, T., et al., Rapamycin increases oxidative metabolism and enhances metabolic flexibility in human cardiac fibroblasts. Geroscience, 2018.

32. Treins, C., et al., Insulin stimulates hypoxia-inducible factor 1 through a phosphatidylinositol 3-kinase/target of rapamycin-dependent signaling pathway. J Biol Chem, 2002. 277(31): p. 27975–81.

33. Maity, A., et al., Epidermal growth factor receptor transcriptionally up-regulates vascular endothelial growth factor expression in human glioblastoma cells via a pathway involving phosphatidylinositol 3’-kinase and distinct from that induced by hypoxia. Cancer Res, 2000. 60(20): p. 5879–86.

34. Miele, C., et al., Insulin and insulin-like growth factor-I induce vascular endothelial growth factor mRNA expression via different signaling pathways. J Biol Chem, 2000. 275(28): p. 21695–702.

35. Masuda, M., et al., A novel frameshift variant of COCH supports the hypothesis that haploinsufficiency is not a cause of autosomal dominant nonsyndromic deafness 9. Biochem Biophys Res Commun, 2016. 469(2): p. 270–4.

36. Bhattacharya, S.K., Focus on molecules: cochlin. Exp Eye Res, 2006. 82(3): p. 355–6.

37. Woulfe, D., et al., Defects in secretion, aggregation, and thrombus formation in platelets from mice lacking Akt2. J Clin Invest, 2004. 113(3): p. 441–50.

38. Pasternak, A.J., et al., Postnatal regulation of MAMDC4 in the porcine intestinal epithelium is influenced by bacterial colonization. Physiol Rep, 2016. 4(21).

39. Wang, Y., et al., miRNA-98-5p Targeting IGF2BP1 Induces Mesenchymal Stem Cell Apoptosis by Modulating PI3K/Akt and p53 in Immune Thrombocytopenia. Mol Ther Nucleic Acids, 2020. 20: p. 764–776.

